# Risk assessment of Ebola virus disease spreading in Uganda using a multilayer temporal network

**DOI:** 10.1101/645598

**Authors:** Mahbubul H Riad, Musa Sekamatte, Felix Ocom, Issa Makumbi, Caterina M Scoglio

## Abstract

Network-based modelling of infectious diseases apply compartmental models on a contact network, which makes the epidemic process crucially dependent on the network structure. For highly contagious diseases such as Ebola virus disease (EVD), the inter-personal contact plays the most vital role in the human to human transmission. Therefore, for accurate representation of the EVD spreading, the contact network needs to resemble the reality. Prior research work has mainly focused on static networks (only permanent contacts) or activity driven networks (only temporal contacts) for Ebola spreading. A comprehensive network for EVD spreading should include both these network structures, as there are always some permanent contacts together with temporal contacts. Therefore, we propose a multilayer temporal network for Uganda, which is at risk of Ebola outbreak from the neighboring Democratic Republic of Congo (DRC) epidemic. The network has a permanent layer representing permanent contacts among individuals within family level, and a data driven temporal network for human movements motivated by cattle trade, fish trade, or general communications. We propose a Gillespie algorithm with the susceptible-infected-recovered (SIR) compartmental model to simulate the evolution of the EVD spreading as well as to evaluate the risk throughout our network. As an example, we applied our method to a multilayer network consisting of 23 districts along different movement routes in Uganda starting from bordering districts of DRC to Kampala. Simulation results shows that some regions are at higher risk of infection, suggesting some focal points for Ebola preparedness and providing direction to inform interventions in the field. Simulation results also shows that decreasing physical contacts as well as increasing preventive measures result in a reduction of chances to develop an outbreak. Overall, the main contribution of this paper lies in the novel method for risk assessment, the accuracy of which can be increased by increasing the amount and the accuracy of the data used to build the network and the model.

## Introduction

Ebola virus disease (EVD) is a viral hemorrhagic fever with a high mortality rate^1, 2^. The virus transmitted via direct contact with blood or other bodily fluids of symptomatic individuals, or with contaminated materials such as bedding, clothing or needles and even direct contact with the dead body of an infected person^2^. People working in the healthcare system at high risk of the EVD. There has been a outbreak of Ebola in the eastern Democratic Republic of Congo (DRC) which is the largest in the history of DRC and the second largest outbreak recorded of Ebola ever (after the 2014-2016 outbreak in West Africa). As of April 26, 2019, 1396 Ebola cases are recorded including 900 deaths and 318 survivors. This is a highly complex Ebola outbreak which is currently transmitting active transmission in 13 of the 21 affected health zones^3^.

The unstable condition due to armed conflict, outbreaks of violence, and other problems in the affected areas complicate public health response activities and increase the risk of disease spread both locally within DRC and to neighboring countries^3^ such as Uganda, Rwanda, Burundi, Zambia, South Sudan, Central Africa etc. With the recent outbreak in DRC, all these countries have the risk of someone infected being entered into and spread the infection there. Therefore, it is very important to public health department to take proper measures to stop entries of infected people into the country. DRC outbreak in neighbouring districts of Uganda posed high risk of Ebola introduction. People from DRC move to Uganda for healthcare, trading and refuge. Therefore, public health department has to prepare proper facilities in case of Ebola introduction. This requires a risk assessment of EVD spreading depending on the human movement as EVD spreads directly via the movement of infected person and their bodily fluid. Modeling the spreading dynamics and risk assessment will enable the public health people to focus their preparedness such as build temporary medical facilities, increasing hospital beds, spread awareness in taking measures such avoid physical contact with infected/unknown people etc. in the high-risk areas.

The availability of accurate models specifically network models for the spreading of infectious diseases and risk assessment has opened a new era in management and containment of epidemics^4–6^. A large number of connectivity driven network models has been proposed for various infectious diseases which are particularly well-suited for capturing the essential features of systems where connections among different nodes in the network are long-lived^7^. A basic assumption with these networks are long-lived contact among individuals indicating links in the networks can be kept constant without oversimplification^8^.

Many models (both network and non-network) for EVD spreading has been formulated assuming a homogeneous long-lived connections among individuals. Different kinds of compartmental models has been used to fit transmission dynamics to the reported data of infected cases for estimating the basic reproductive ratio^9–12^, international spreading risk^13^, to suggest mitigation measures^14–16^.

However, long-lived connections not true for systems where links are rapidly changing^8, 17^. Highly contagious/infectious diseases are mostly transmitted form an infected individual to susceptible via physical contact^18^. Therefore, models for these contagious disease spreading are crucially dependent on the contact structure among individuals (nodes) in the network^19^. If we consider a network models which consists of human, then the assumption of their contact with each other being constant is actually an oversimplification of the reality. In general, the contact structure among individuals change with time. However, these changes in contacts are not completely random, rather there are always some patterns. For example, the contact pattern is will change based on the change of frequency of the occasional or permanent partners in case of sexually transmitted disease. Therefore, the change of the permanent partners can be a long term process and for that, connectivity driven network models work fine. However, when we consider the occasional partner change, this connectivity driven network will not be able to capture the frequent change of partners. To adapt notwork models to this changing contact pattern, several approaches has been proposed over time. One of the earliest concept is the switching networks^20, 21^. In this model, the contact network switches among some predetermined networks structures. This model takes into account the changing structures of the network with time. However, we have to define some predefined structure for this type of network model. Another approach is activity driven network, which has the ability to capture the dynamic contact pattern in the network^7^. Activity driven network has been greatly used to capture the transmission dynamics of infectious diseases such as Ebola spreading. In these activity driven network, an activity firing rate is applied to the individuals and according to that rate, each node is activated^7^. After the activation, that node will have the ability to connect with a predefined number of nodes in the network.

Activity driven networks (ADN) has been widely used for EVD spreading. ADN overcomes the simplifying assumption of long-lived and homogeneous contacts among individuals widely used in previous compartmental models^6^. Rizzou et el. used ADN to emulate the dynamics of EVD in Liberia and offer a one-year prediction^6^. The effect on contact tracing on the spreading dynamics has also been quantified using ADN^22, 23^

However, this activity driven network has a limitation as it randomly creates new links at every time steps. Therefore, if there are any permanent links in the network, that is not considered in the activity driven network and it can be a problem for ADN. Vajdi et al. proposed a temporal network to overcome this problem with the ADN^24^. This multilayer temporal network incorporates both permanent links and occasional links in two different layers.

In this paper, we develop a novel method for risk assessment using a multilayer temporal network, a modified SIR model for temporal network named susceptible active-susceptible inactive-infected active-infected inactive-recovered (*S*_*a*_ *S*_*i*_ *I*_*a*_ *I*_*i*_R) model, and the Gillespie algorithm. The multilayer network has two layers— a permanent layer reflecting permanent contacts and a temporal layer that incorporates potential contacts. We adapt Gillespie algorithm with the *S*_*a*_ *S*_*i*_ *I*_*a*_ *I*_*i*_ R compartmental model and the multilayer network to see the evolution of disease spread and risk assessment.

As an example of the method, we propose a multilayer temporal network for Uganda including 23 districts, based on human movements from a focal bordering district of DRC to Kampala where permanent layer expresses contacts among family members while intra- and inter-districts contacts reflects potential contacts due to human movement. Simulation results suggest that making people aware to reduce physical contact while travelling and taking other preventive measures will reduce number of EVD infected human. Results show that some districts are more vulnerable to the risk of EVD spreading than others, which may suggest important guidelines for public health personnel in applying interventions and prioritizing resource allocation. Assessed risks are probabilities of infection spreading for our specific scenario based on generic and incomplete movements data and any change in the network will results in different risks. Therefore, risk assessments in this paper are just some examples of our proposed novel risk assessment method. The main contribution of this paper is the novel multilayer temporal network based simulation tool for risk assessment of EVD spreading, which has the capability to be used in practical purposes when incorporating accurate movement data and model parameters. The rest of the paper is organized as follows. The Risk assessment method section describes the multilayer temporal network, *S*_*a*_ *S*_*i*_ *I*_*a*_ *I*_*i*_R epidemics on multilayer temporal network and the adaptation of Gillespie algorithm for risk assessment. Application of risk assesment for Uganda EVD spreading showed an example for multilayer temporal network in Uganda, simulation results and discussion. We summarize our conclusions and suggestions drawn from simulation results in the Conclusion section.

## Risk assessment method

In this section we propose the novel method for risk assessment using multilayer temporal network, *S*_*a*_ *S*_*i*_ *I*_*a*_ *I*_*i*_R spreading model and the adaptation of Gillespie algorithm for temporal network.

### Multilayer temporal network

We consider a population of *N* individuals connected with two different types of links. These individuals can have links in two different layers. In the first layer *L*_1_, the links are considered permanent. Therefore, in this layer, contacts among individuals are not changing. There are potential links in the second layer *L*_2_ based on some certain probability distributions. These potential links/contacts are activated once individuals at the both end of this link are active simultaneously. The activation of the individuals are driven by a activity rate (*γ*). The intersection of the two network layers are assumed empty. Once both nodes are active, the link between them become active with a probability *P*_0_. This probability can have a different value depending on the nodes the link is connecting. Therefore, this temporal network has the ability to incorporate the heterogeneous contact probability. However, in this work a single *P*_0_ is used in the absence of specific information about the contact between active node pairs. In the subsequent section of this work, we call *L*_1_ as permanent layer while *L*_2_ is called temporal/potential layer^24^. The generalized structure of multilayer temporal network is in presented in Figure 1.

**Figure 1.**
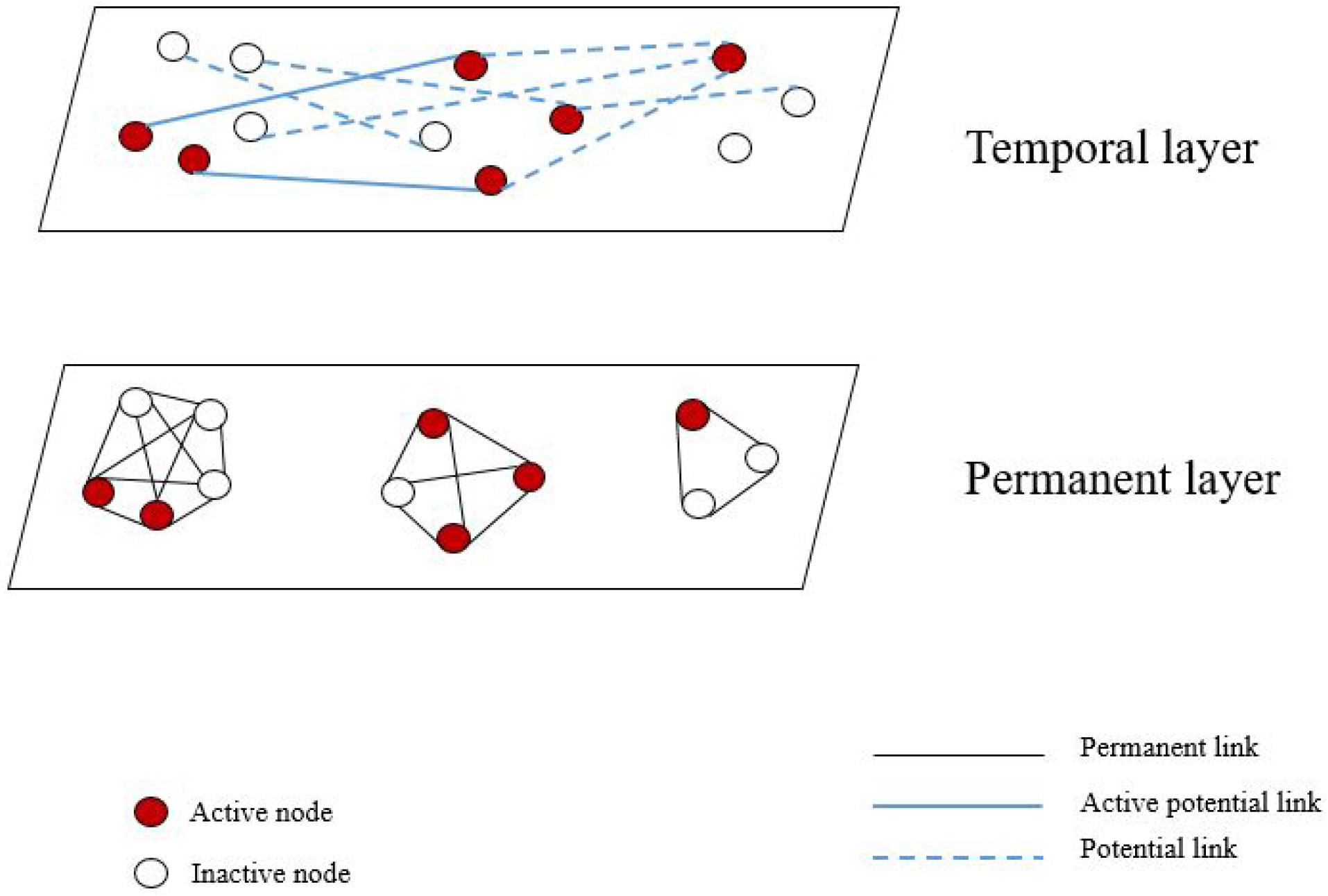
A generalized representation of the multilayer temporal model at a specific time *t*. White circles represents inactive nodes while red circles represents active nodes. Separate rectangles represent two different layers. Dark solid lines show permanent links while coloured lines show links in the potential layer. A Potential link becomes active with a probability *P*_0_ when both ends of the link are active nodes.

In the permanent layer, a link can always transmit infection irrespective of its active status while a link in potential layer transmits the infection only when it connects two active individuals. At a single time step, one node can only be active or inactive. When a node becomes active, it can come in contact with the active potential active neighbours in *L*_2_ and with a probability *P*_0_, it activates the potential contact. When one of the nodes in the in the contact becomes deactivated, the link vanishes and the probability of infection through that link becomes zero again. We assume this process of a node being active or inactive a Poisson process where a node becomes active and deactivated with rate *γ*_1_ and *γ*_2_ respectively. Therefore, a node will stay active for a period of 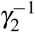 as an active node changes its state with a rate *γ*_2_. Thus we can assume a high value of *γ*_2_ and *γ*_1_ for a specific node will reflect the reduced duration of the potential link. In this work we assume a similar value for therefore in the subsequent section, we express *γ*_1_ and *γ*_2_ as *γ* only and refer it as activity rate. Therefore, increased *γ* means an increasing frequency of a node becoming active and inactive. For example, if we assume an individual becomes active once he starts a movement/trip and stays active until the trip is finished then a decreased *γ* means an length of the trip and an increase in the frequency of this kind of trips. Moreover, if a node does not participate in the occasional contacts —it never becomes active— *γ* is set equal to zero for that node. Inactivation times for nodes are exponentially distributed, and therefore the temporal contact duration has an exponential distribution. In fact, a temporal contact disappears with the inactivity of a node or both in the link^24^. This temporal network is different from the widely used activity driven network in the contact structure among individuals. In the activity driven network, links are created and destroyed completely randomly and there are no permanent links. In the next time period, all these links are removed and new links are created with a specific node when it becomes active.

### *S*_*a*_*S*_*i*_*I*_*a*_*I*_*i*_R epidemics on multilayer temporal network

In this section, we describe the modification of Susceptible-Infected-Removed (SIR) model for our multilayer temporal network. SIR model is a popular approach for studying infection spreading where infected people dies or eventually recovers and gains life-long immunity. For disease such such as Chicken pox, EVD etc., SIR model has the potential to describe the disease dynamics and infection spreading. In SIR model, each individual is either susceptible, infected, or removed/recovered. We assume that the infection and recovery processes are independent Poisson processes. A susceptible node becomes infected with EVD if it comes in contact with an infected person and this transition happens with a rate *β*. This is called transmission rate. Once a person becomes infected, he/she stays infected for a certain period of time. This period is called infectious period. After the infectious period, individuals either recovers/removed from the infection. The rate at which an infected person leaves the infected state in called recovery/removal rate and it is inverse of the duration of the infection. For our SIR model, the recovery/removal rate is denoted as *δ* which has a unit *time*^−1^. So far we have discussed the basic SIR model. However, for our multilayer temporal network, we have also status of the individuals (active/inactive). Combining our temporal network model and SIR spreading process, we can have total five states that an individual can occupy. Therefore, we named the model as *S*_*a*_ *S*_*i*_ *I*_*a*_ *I*_*i*_R model, where compartments are inactive susceptible, active susceptible, inactive infected, active infected and recovered.

If the probability of node *i* occupying inactive susceptible, active susceptible, inactive infected, active infected and recovered in the mean-filed approximation are expressed as *S*, *S*_*a*_, *I*, *I*_*a*_, and *R* respectively, then equations for the time evolution can be expressed as-

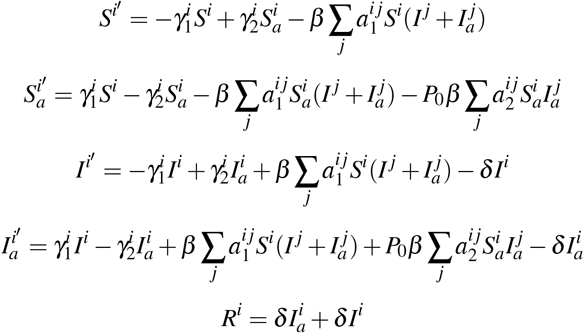

Where *P*_0_ is the probability of a potential link becoming active once both end of the link is active. 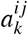 is the element of adjacency matrix *A*_*k*_ where *k*=1 for permanent layer and *k*=2 for potential layer.

### Adaptation of Gillespie Algorithm

Gillespie algorithm has been widely used to simulate stochastic processes for static network (permanent contacts)^25–29^ and dynamic/time-varying networks (temporal/potential contacts)^24, 30^. However, our multilayer temporal network has both static and temporal contacts. Therefore, Gillespie algorithm was adapted to the changed network state at every time step retaining permanent contacts.

- Initialize the number nodes in the network (*N*), state transition matrices, maximum number of events and the final time for simulation. Find the transition probability *R*_*i*_ for each node at the next time step and *R*_*tot*_ = ∑_*j*_ *R*_*j*_, where *j* = 1, 2,…*N* and keep track of the status of the nodes (active/inactive)
- Find the time for the next event which is exponentially distributed with a rate *R*_*tot*_. The second step is to select a node according to the probability distribution *Pr*(*i*) = *R*_*i*_/*R*_*tot*_ which will make a state transition. Later we select a state where the transition will happen.
- Increase the time by time calculated in Step 2. The main difference lies in the update of transition rates for each nodes in the network once a transition occurs for our adaptation of Gillespie algorithm. Nodes can change their status (active/inactive) as well as changing their states (susceptible, infected, recovered). Therefore, every time a transition occurs, the status of the nodes needs to updated and recorded. When a node is in the permanent layer, it’s transition probability has the impact of both layer’s neighbours. However, when the node is in temporal layer, it only has the impact of its neighbours on the temporal layer. Therefore, the algorithm is modified to account for the network with two different layer’s impact on the transition probabilities.
- Go back to Step 2 unless the stop condition is reached (maximum number of events, the final time for simulation, or *R*_*tot*_ < *Tolerances*).

### Calculation of Risk

As the spreading process in our multilayer-based temporal network was highly stochastic, we performed two hundred simulations for each combination of parameters. We kept track of each node’s status and counted the numbers of simulations in which a particular node was infected. This count was later used to calculate the risk of EVD spreading in each district. We used the following formula to calculate the spreading risk to a specific district–

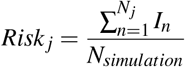

Where, *Risk*_*j*_ = Ebola spreading risk of district *j*, *I*_*n*_ = Number of simulations where node *n* is infected, *N*_*simulation*_= Number of simulations, *N*_*j*_= Total number of population in *j*^*th*^ district, and *j* = 1, 2, 3,…, 23.

Once we calculate the risk, we can use any mapping software such as ArcGIS software to create risk maps. Risk maps provide a visual representation of of spreading risks.

## Application of Risk assessment for Uganda EVD spreading

We applied our risk assessment method for EVD spreading using generalized movement pattern and a some specific model parameters. The application of our developed method is presented in this section. First we formulate the multilayer temporal network for Uganda. Later we used the network with *S*_*a*_ *S*_*i*_ *I*_*a*_ *I*_*i*_R model and the modified Gillespie algorithm to track number of infected human after a certain period of time and assess the risk of EVD spreading to different spatial location. Simulation results and discussion are presented for specific scenarios are presented at the end of this section.

### Multilayer temporal network for Uganda

We have proposed a multilayer temporal network for Ebola spreading in Uganda. The recent Ebola outbreak in Democratic Republic of Congo created a possibility of EVD infection in Uganda. We have created a network based on the people’s movement for different purposes such as fish trade, cattle trade and general movement for different purposes. The movement pattern and districts in the movement paths are obtained from confidential data provided by Ministry of Health, Uganda. These data are then used to formulate an example of human movement network for some selected districts in Uganda. We have observed the focus of Ebola preparedness by the Ministry of Health and partners to select possible point of entry for Ebola patient from DRC. We found out that Kasese district is at the high risk of EVD infected person’s entry point. Therefore we choose this district as our point of entry. Once an infected person from DRC enters into Uganda through the bordering districts, they come in contact with susceptible people in that location. Now, Ebola being highly contagious, infected person/persons can spread the infection to people they come in contact with. People moves from one location to another for different reasons and there is always a pattern for this movement. For Uganda, we have obtained some general information about the local peoples movement pattern and incorporate them in our multilayer network. People movement from one location to another are mostly motivated by several reasons. For Uganda, we found out that these movements are mostly for three different purposes. They are following-

1. Fish trade towards southward direction from the point of entry
2. General movement for daily commute, shopping, visiting relatives, search of work, or travelling for various purposes. These types of movement starts in the point of entry in the bordering districts and goes all the way to capital city Kampala. People mostly travels from rural areas to neighbouring big cities. Once they come in contact with other people there and this results in a certain mixing among individuals from locations. This mixing happens throughout this movement path. There are several of this movement paths which are presented in the Figure 2 with green arrows.
3. Limited movements due to cattle trade. The movements for cattle trade are mostly local or between neighbouring districts. long distance cattle trade happens in commercial scale and does not include much movement of people as they happens mostly via organized transportation system.

**Figure 2.**
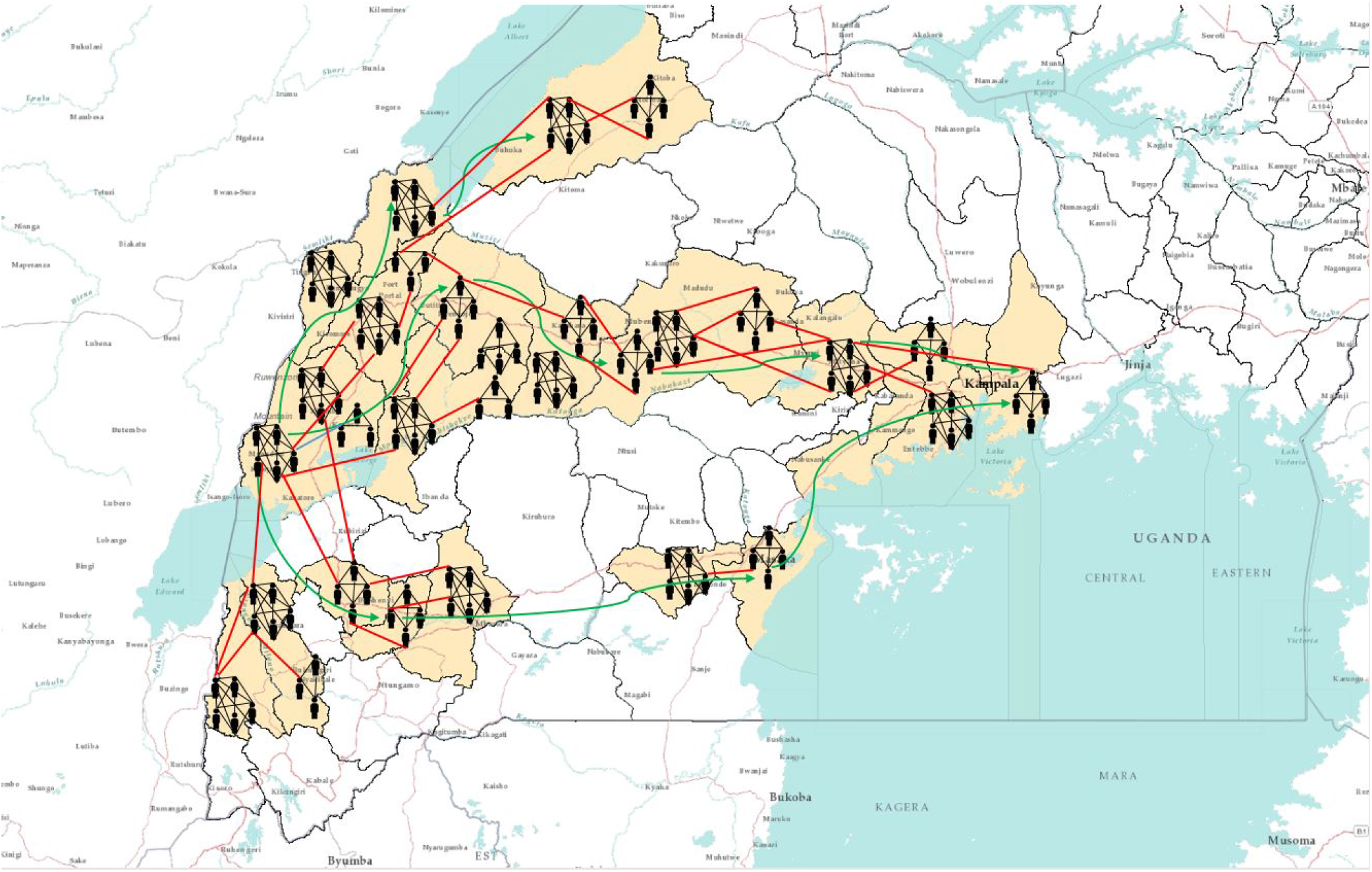
Multilayer temporal network for EVD spreading in Uganda. The districts considered for our network model are coloured in baize (23 districts). Small human shapes represents individual human beings. This figure does not represent the actual network, rather it is a visual representation of the multilayer network. Each cluster of human represents a households and black lines among them represents permanents contacts. Red lines represent contacts in potential layer. These links only become active when both individuals in the link are active simultaneously. A link represents a possibility of pathogen transmission. Green directional arrow represents the directions of human movements.

To represent this contact structure and movements in Uganda, we propose a multilayer contact network. We create a specific network for human movement from Kasese to Kampala based on general movement information. The network is used to show an example of the developed method for risk assessment. As we have described in our Temporal network section, we assumed two layers in the network. The permanent layer consists of the contacts of each individuals within each household. Each family in Uganda has an average of 6 children^31^. Therefore, with the parents we can assume around 8 people in a household in average. We assume a distribution of population where each household can have 5-10 members.

Within each household, we can assume permanent links among family members. Therefore, links within each household can construct the permanent layer in the Uganda network for Ebola spreading. The potential layer can be formed incorporating individuals movement for different purposes. An individual becomes active once it is in the movement and stays active until it finishes its movement. Once a node is in movement (active), its potential links with individuals other than own family can be activated based on the active status of the other nodes that has link with our specific node. This can be explained as following-node *i* has potential links with a set of node named *J* throughout the whole network. Therefore, once node *i* is active, potential links with any of the node in *J* can be activated if that node itself is active as well. This link activation structure is very crucial in the proper representation of the contact structure. As both of these node are in movement, they can come in contact with each other in different places such as transportation, marketplace, visiting sites etc. Usually the movement pattern between individuals follow a general structure. Within each location, there is a probability that individuals will come in contact with each other for various purposes. Inter-location contacts also follow some structure rather than being completely randomized. People usually flock into big cities or towns nearby where they will come in contact with the local active people or others coming in to that location for similar purposes. When this happen, if two of these active individuals have a potential link, that will be activated and there will be a possibility of infection via that link.

Districts we have used in our Uganda network are presented in Table 1. We have used the centroid of each district to find the distance between two districts while formulating the network. However, this is one realization of the network built using available movement data. Some Ugandan districts are not included in our assessment, therefore there may have been districts at high risk that are excluded from the risk assessment performed in this article. We have scaled the population in each district by 1000 in our network for computational purposes. As our main purpose is to evaluate the risk of EVD, this scaling has greatly reduced the computational complexity. We have incorporated in the network all previously discussed movement patterns along different paths from one point of entry to Kampala. These movements and their directions are also shown in Figure 2. We have considered all districts that are in movements paths of any nature towards Kampala from Kasese. Although all bordering districts are at risk of Ebola introduction, we focus on demonstrating the application of our method when the initial infections are in the Kasese district.

**Table 1.** Districts considered in our Uganda multilayer temporal network

| Districts | Population |
| --- | --- |
| Bundibugyo | 224387 |
| Bunyangabu | 181200 |
| Bushenyi | 234440 |
| Hoima | 572986 |
| Kabarole | 469236 |
| Kampala | 1507080 |
| Kamwenge | 414454 |
| Kanungu | 252144 |
| Kassanda | 100038 |
| Kasese | 694992 |
| Kyegegwa | 159800 |
| Kyenjojo | 422204 |
| Lwengo | 267300 |
| Masaka | 297004 |
| Mbarara | 472629 |
| Mityana | 328964 |
| Mpigi | 250548 |
| Mubende | 684337 |
| Mukono | 594804 |
| Ntorko | 70900 |
| Rukungiri | 314694 |
| Sheema | 180200 |
| Wakiso | 1997418 |

In summary, our network for Ebola spreading in Uganda consists of 23 districts where permanent layer incorporates the contacts among individuals within a household. Potential layer reflects the contacts between individuals which can happen only when both ends of the link are active due to some kind of movement and they have a possibility of pathogen transfer between them.

### Simulation Setup

Upon formulation of the multilayer network, we perform simulations with the *S*_*i*_ *I*_*a*_ *I*_*i*_R model for EVD spreading using Gillespie algorithm^24, 32^. The network is temporal as depending on the active status of nodes, links are changing in the potential layer. We conduct our simulation with two major goals— observe the progression of EVD spreading with time throughout the network and evaluate the risk of spatial spreading for this specific scenario.

In our temporal simulations, we have four different parameters. They are the rate at which individuals become active and inactive (*γ*_1_ and *γ*_2_, generally we will call them activity rate *γ* as both of them have same value for our simulations), the transmission rate *β*, the recovery rate *δ*, and the probability *P*_0_ of pathogen transmission via a potential link between two simultaneously active individuals in the potential layer. These values are hard to estimate and due to high stochasticity in the people’s movement pattern, movement parameters i.e. (*γ*) can not be expressed with a single value. The transmission rate *β* is also variable and takes different values for different outbreaks depending on the contact pattern among individuals. Therefore, we perform simulations with multiple values of each parameter to explore the sensitivity of the epidemic size and and spreading risk to different districts from the initial outbreak location. For each parameter set, we calculate total number of infected individuals with time and create risk map.

We present the number of infected individuals in each time step for each parameter set. As we have multiple parameters, we choose the value of *P*_0_=0.7 and 0.1 which represents the seventy percent and ten percent chance of the pathogen transmission via a potential link between two active nodes. For the value of rate a node becomes active and inactive, we choose *γ*= 0.1 and 0.5. This can be explained as following, a particular individual becomes active in every 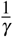 days and once active, it stays active for next 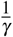 days. For example, *γ*= 0.1 represents the frequency of a node being changing its state (inactive to active, active to inactive) in every 10 days.

The value of the transmission rate is mostly estimated from the real incidence data in most of the literature where Ebola model is fit to the past incidence data. For our work, we are more interested in the risk assessment and provide different scenarios under different human movement and weather conditions. Therefore, a range of parameter values are used to quantify the size of the epidemic and assess the risk of EVD spreading to different locations from the initial outbreak location in Uganda.

## Results and Discussion

We initiated our simulation with a single active infected person in Kasese district and tracked the status of each node in the network for a period of 150 days to see how each node is changing their status (active susceptible, inactive susceptible, active infected, inactive infected, and recovered). At the end of the simulation time (150 days), total number of infected people in the outbreak are calculated and the risk of a specific node being infected during the outbreak was assessed.

To measure the progression and the severity of Ebola spreading, we track total number of infected and cumulative number of infected people for a period of 150 days. For any infection spreading, some important measures are the size of the peak infection (maximum number of simultaneously infected individuals), time to reach that peak and total number of infected within an outbreak (epidemic size)^29^. Therefore, we designed our simulation to keep track of both the number of infected and cumulative number of infected at each time step of 150 days simulation period. Our simulation results are presented in the subsequent part of this section. We have chosen values of transmission rate *β* to explore varying range of transmission potential given a contact with infected individual. We have presented simulation results for *β* =0.2, 0.5, 1.7, 2.5 in the results section.

Figure 3 represents the number of cumulative infected and infected humans for *P*_0_ =0.7 and *γ*=0.5 and a varying range of transmission rate *β*.

**Figure 3.**
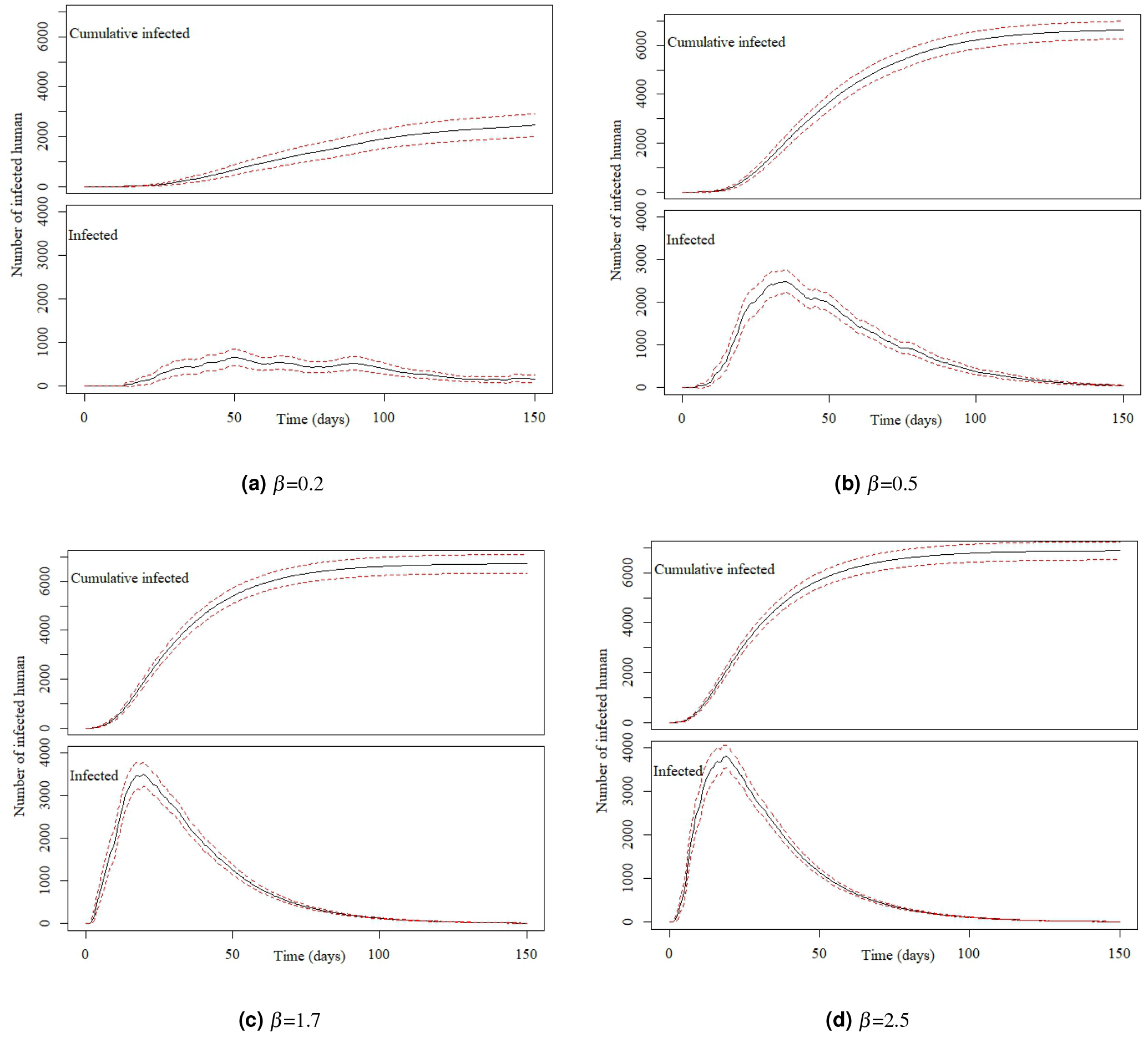
Number of cumulative infected and infected individuals in the Uganda Ebola network with 95% confidence interval for *P*_0_=0.7, *γ*=0.5, and, (a) *β* =0.2, (b) *β* = 0.5, (c) *β* =1.7, and (d) *β* = 2.5

Within each sub-figure in Figure 3, the top graph represents the cumulative number of infected while the bottom plot represents number of simultaneously infected humans at different time steps.

From Figure 3 (*P*_0_=0.7 and *γ*=0.5), it is evident that with the increase of *β*, the infection size increases rapidly. An increase of *β* from 0.2 to 0.5 increases the infection size from 2459 to 6634. Therefore, very small increase in the transmission causes a huge increase in the total number people infected with Ebola when we assume a 70% chance of pathogen transmission once an active infected individual comes in contact with another active susceptible person in the potential layer. Also, in this simulation set, the potential layer assumed to be highly active/mobile as people assumed to become active and inactive within a period of two days.

The number of simultaneously infected people at a certain time is very important for public health personnel^33^. More infected people means preparation of more hospital beds, doctors and medical supplies^34^. Therefore, once a outbreak occurs, it is very important to have an idea about the maximum number of simultaneously infected people (size of the peak infection)^29^. In each of the sub-figure, the bottom plot shows simultaneously infected people at each time step. In the plot, the peak infection size also increases with *β*. However, when *β* =0.2, the peak is not very pronounced and peak infection size is around 600. However, for other values of *β*, The peak is more pronounced and the peak infection size is more than 2500 for all other values of *β*. The time to reach the peak for for *β* =0.2 is around 50 days. With the increase of *β*, the the infection plot becomes skewer to the left, meaning faster arrival to the peak infection size. Therefore, faster arrival to peak infection and greater value of peak infection size indicates a widespread epidemic outbreak. Therefore, simulations for this specific parameter set indicates a widespread and severe outbreak for for *β*>0.5 where more than 50% of our total population in the network becomes infected.

For each parameter set, we have estimated the risk of EVD spreading to different spatial location when we start the infection at Kassese district with a single infected individual. We have created risk maps using ArcGIS for the spreading of Ebola to distant locations. We have used equation for Risk explained the “Material and Methods” section to calculate the risk. Value of the risk is classifies in 5 different categories as presented in Table 1.

Risk maps in Figures 4, 6, 8, and 10 show all the neighboring districts in the south part of Uganda although we have considered only 23 districts, which are considered being in the path of people’s movement network from Kasese to Kampala. Therefore, there are some districts in the maps that have not been considered in our multilayer temporal network and hence have not been assessed for risk.

**Figure 4.**
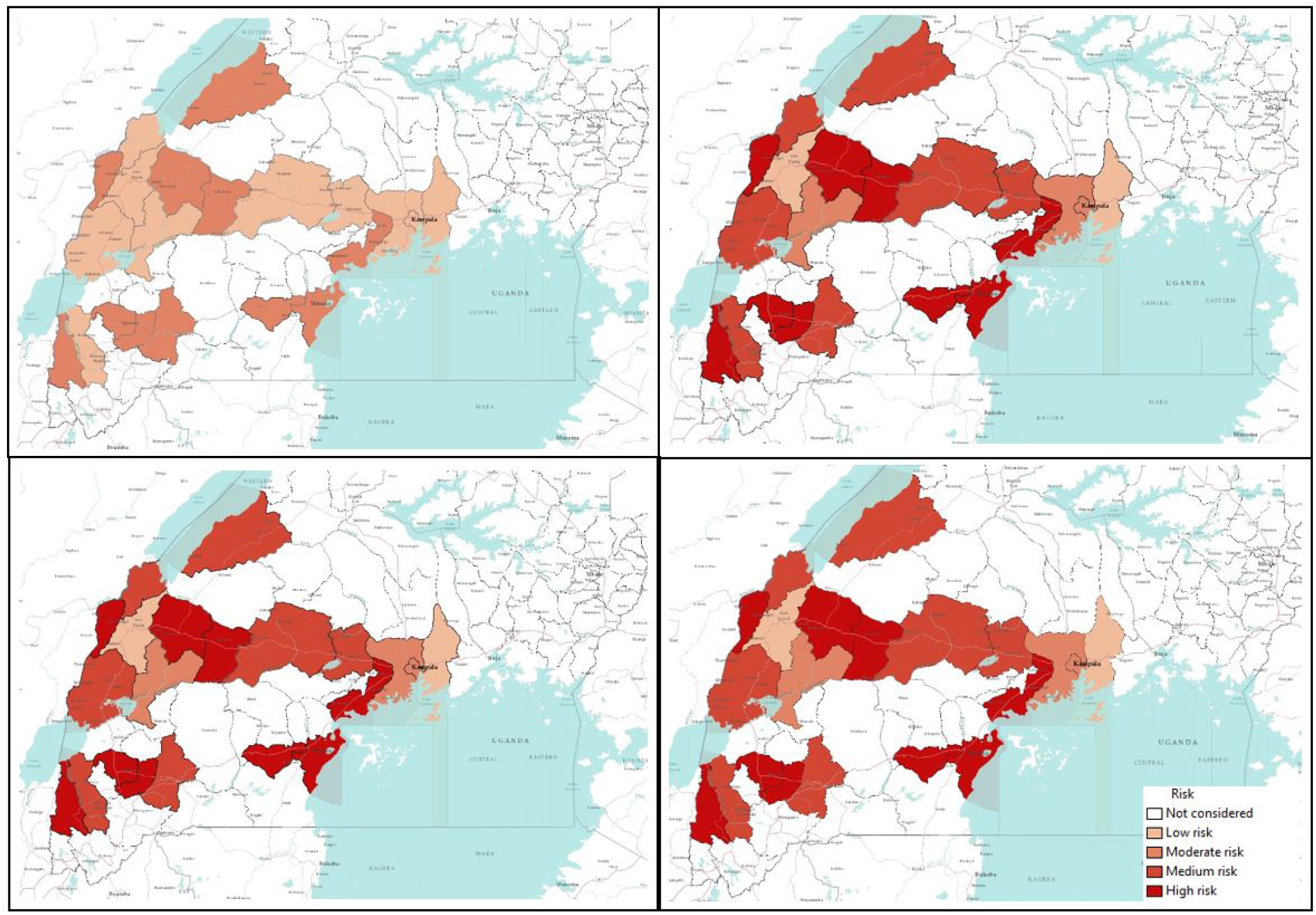
Risk map of Ebola spreading within selected 23 districts in Uganda for *β* =0.2 (top left), 0.5 (top right), 1.7 (bottom left), 2.5 (bottom right). The map is colour coded according to the risk of Ebola spreading.

Figure 4 shows a risk map for four selected values of *β* and *P*_0_= 0.7 and *γ* = 0.5. The map is coloured with a monochromatic colour gradient which increases with increase of risk. Therefore, districts which are white in the map are either not at risk (if considered in the network model) or not considered in the network model.

From the risk map for *β* =0.2 in Figure 4, we can see that all our selected locations are at low risk or moderate risk as the transmission rate is very low. However, some of the districts are at comparatively high risk than others. For this specific scenario with *β* =0.2, Bundibugyo, Bushenyi, Kyegegwa, Kyenjojo, Masaka, Mpigi, and Sheema districts are at higher risk than other districts in our network. However, aforementioned districts are at moderate risk of Ebola spreading while other districts are at low risk for *β* =0.2. Increasing the value of *β* increases the risks proportionately which can be seen from Figure 4. The colour gradients are increasing in the risk maps with the increasing *β*. In the risk maps for other values of *β*, a similarity in the risk is observed. This is evident from similar colour gradient of districts in all three maps for *β* =0.5, 1.7, and 2.5, which can be explained from similar epidemic size and comparable infection peaks for these values of *β* (Figure 3). Therefore, we have a similar infection spreading in these cases and assessed risks are similar (not same, but they are in the same interval presented in Table 1). There was an increasing trend in the value of calculated risk parameters with the increase of *β*, although in the map they are included in the same interval. Districts in the temporal network for *β* = 0.5, 1.7 and 2.5 with assessed risks are presented in Table 2.

**Table 2.** Classification of risk for our spatial locations based on the value of risk parameter.

| Risk | Value of risk parameter |
| --- | --- |
| No risk or not considered | 0 |
| Low risk | $0 < \text{risk} \leq 0.2$ |
| Moderate risk | $0.2 < \text{risk} \leq 0.4$ |
| Medium risk | $0.4 < \text{risk} \leq 0.6$ |
| High risk | $> 0.6$ |

Table 3 shows that 10 of 23 districts are at high risk of EVD spreading in case of an outbreak. With current outbreak in DRC, it is expected that bordering districts will be at high risk of EVD spreading. However, our simulation results has the capability to incorporate the movement data that shows non-bordering districts can also be at high risk due to infected person’s movement. This is can be easily seen from Figure 4, where some non-bordering districts demonstrates high or medium risk of EVD spreading.

**Table 3.** Districts in the network and their associated risk for EVD spreading

| Risk | Districts |
| --- | --- |
| High risk | Bundibugyo, Bunyangabu, Bushenyi, Kanungu, Kyegegwa, Kyenjojo, Lwengo, Masaka, Mpigi, Sheema |
| Medium Risk | Hoima, Kasese, Mbarara, Mityana, Mubende, Ntoroko, Rukungiri, Kassanda |
| Moderate risk | Kampala, Kamwenge, Wakiso |
| Low risk | Kabarole, Mukono |

Figure 5 represents the number of infected individuals for *P*_0_ =0.7 and *γ*=0.1. *γ* = 0.1 means individuals have the probability of becoming active and inactive in every 10 days. Therefore, decreasing the *γ* decrease the probability of a human becoming active while increasing the time of him/her staying active.

**Figure 5.**
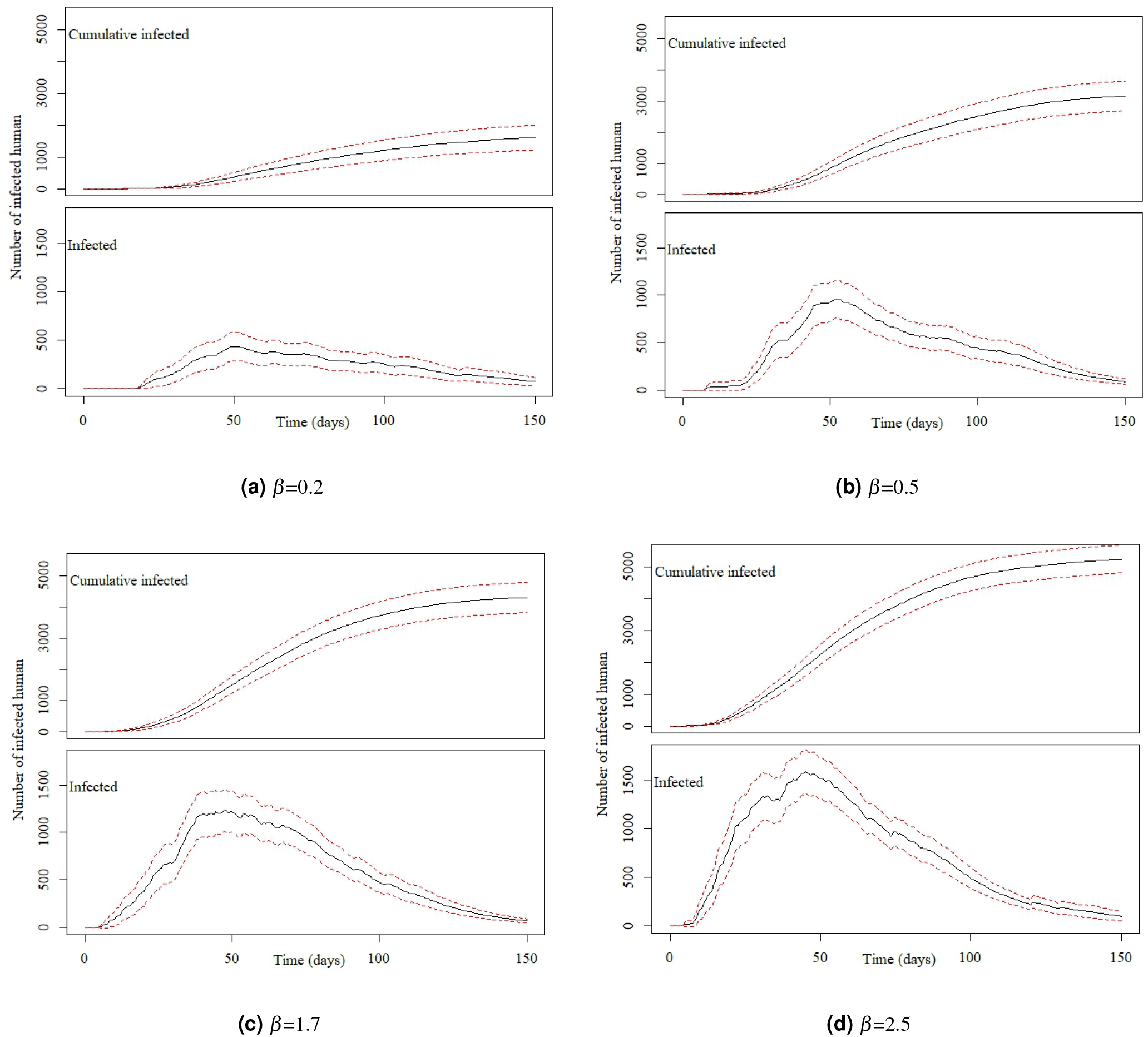
Number of infected individuals in the Uganda Ebola network with 95% confidence interval with *P*_0_=0.7, *γ*=0.1, and for (a) *β* =0.2, (b) *β* =0.5, (c) *β* =1.7, and (d) *β* = 2.5

The peak infection increases from 450 to 1700 for our selected values of *β* =0.2 to 2.5 while the cumulative number of infected increases from 1607 to 59.

Comparing the simulation results in Figure 3 and 5 for similar value of *β*, the epidemic size as well as the peak infection is always higher for higher values of *γ*. Higher value of *γ* can be translated to the increased human movement in our network. As we are assuming that a human is active when it is in a movement, a higher *γ* for a person means frequent short trips while lower *γ* means longer trips which are not very frequent.

Higher *γ* increases the time a human will be in the active (mobile) state, While lower value indicates longer length of an individual staying active^24^. Therefore, it is convenient to assume that while an infected individual is active for a longer period of time, it will eventually spread the infection to more active individuals in the potential layer^35^. However, our simulation results show otherwise. An increase in the value of *γ* increases the infection size as well as the peak infection size (Figure 3 and Figure 5).

Therefore, our simulation results show the frequency at which individuals become active or inactive dominates over the time span of the individual staying active. This is evident from the higher size of the epidemic from the higher value of *γ*. For *β* =0.2, the cumulative number of infected after 1607 when it is 59 when we have *γ*=0.5. The value of the cumulative number of infected is also higher for other cases as well. The infection reaches to the peak infection slowly (approximately 50 days) irrespective of the value of the transmission rate. Therefore, lower value of the activity rate *γ* means lesser number of simultaneously infected people and slow infection spreading within the spatial locations (Figure 4 and Figure 6). As we have discussed earlier, lower value of *γ* reflects the reduced movement of people. Therefore, from the comparison of simulation results in Figure 3 and Figure 5, it is evident that human movements are critical in the severity and speed of EVD spreading. As frequent human movements (frequent short trips) spreads the EVD very fast, reduced human movement may minimize the severity of the EVD spread^36^. This is also evident from the risk map showed in Figure 6 for reduced *γ*. From the map, we can see that Bundibugiyo, Sheema and Masaka are three districts which are most at risk of Ebola spreading for our specific network model. For these three districts, the risk of spreading is comparatively higher than other districts even when the value of the transmission rate is very low. For example, these districts are in moderate risk zone while others in low risk zone for *β* =0.2. However, with the increase of *β*, the value increases and for our highest value of *β* =2.5, these three districts are at the highest risk of Ebola spreading. Table 3 presents districts in the temporal network for *β* = 2.5 with assessed risks.

**Figure 6.**
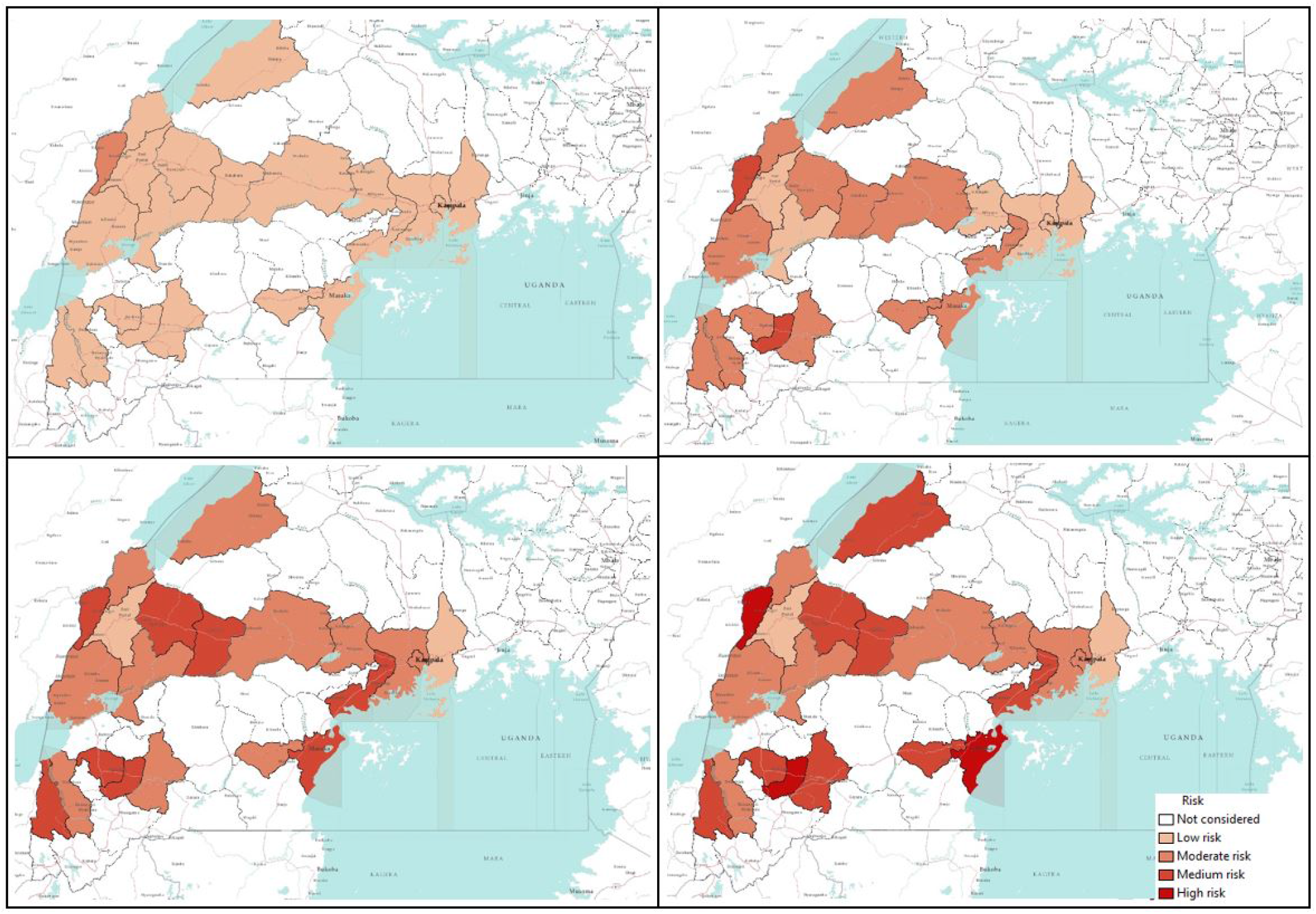
Risk map of Ebola spreading within selected 23 districts in Uganda for *β* =0.2 (top left), 0.5 (top right), 1.7 (bottom left), 2.5 (bottom right). The map is colour coded according to the risk of Ebola spreading.

During the network creation, some of the districts were assumed to be possible mixing places, which makes them more vulnerable than others. Therefore, simulation results and evaluated risks are obtained according to the network structure. An example of dependency on the network structure is evident from Figures 4 and 6, showing the district of Kabarole at a low risk for all values of *β*, despite being a bordering district to DRC. This low risk for Kabarole can be attributed to the fact that this district was not considered as a mixing place in our network. This demonstrates our method’s adaptability to specific data about each locations in the network.

**Table 4.** Districts in the network and their associated risk for Ebola spreading for *P*_0_=0.7, *γ*=0.1 and *β* =2.5

| Risk | Districts |
| --- | --- |
| High risk | Bundibugyo, Masaka, Sheema |
| Medium Risk | Bunyangabu, Bushenyi, Hoima, Mbarara, Mityana, Mpigi, Kyegegwa, Kyenjojo, Lwengo, Kanungu |
| Moderate risk | Kampala, Kamwenge, Wakiso, Rukungiri, Ntoroko, Mubende, Mityana, Kasese, Kassanda |
| Low risk | Kabarole, Mukono |

Comparing table 2 and Table 3, it is evident that when we have a lower *γ*, risks decreases significantly. For same values of *β*, lower *γ* reduce the risk of Ebola spreading. When *β* ≥ 0.5 and *γ*=0.5, there are eleven high risk districts. Decreasing the value of *γ* to 0.1 results in only 3 high risk districts for *β* =2.5. For other values of *β* and *γ*=0.1, none of the districts in our network model are at high risk. Therefore, reducing human movements has shown significant decrease in the Ebola spreading. However, reducing human movement is not practical as human movements can not be controlled. Therefore, we focus on the parameter *P*_0_ which express the possibility of pathogen transfer via an active potential link. *P*_0_ can be expressed as the probability of an active infected person spreading the virus to an active susceptible person. The value of this parameter is dependent on direct physical contact between infected and susceptible. In our previous set of simulations, we used *P*_0_=0.7 in our previous simulations. Now if we can decrease the possibility of the physical contacts among humans in case of an outbreak, it would be equivalent to a reduced possibility of virus transfer. To observe the impact of reduced physical contact i.e. lower *P*_0_, we conducted simulation for a *P*_0_=0.1 reflecting only 10% possibility of pathogen transfer via an active link while one of the human is infected. Simulations results for *P*_0_=0.1 are presented in Figures 6-9.

Decreasing the probability of Ebola spreading (i.e. *P*_0_) to 10% from 70% via a contact in potential layer significantly reduces the infection size as well as simultaneously infected humans. However, while changing the *P*_0_, similar values of *γ* are used as before.

Figure 7 shows cumulative infected and infected humans for *γ*=0.5 and *P*_0_=0.1. For our lowest value of *β* =0.2, cumulative number of infected human are 45 after 150 days. Increasing *β* increased the value of cumulative infected humans to 1274 for *β* =2.5. Therefore, number of cumulative infected humans are very low compared to cumulative infected humans for similar values of *β* with *P*_0_=0.7. Also there is no pronounced single peak while maximum simultaneously infected people goes around 400 for the highest used value of *β*. Therefore, decreasing *P*_0_ reduces number of infected humans and therefor the severity of EVD spreading.

**Figure 7.**
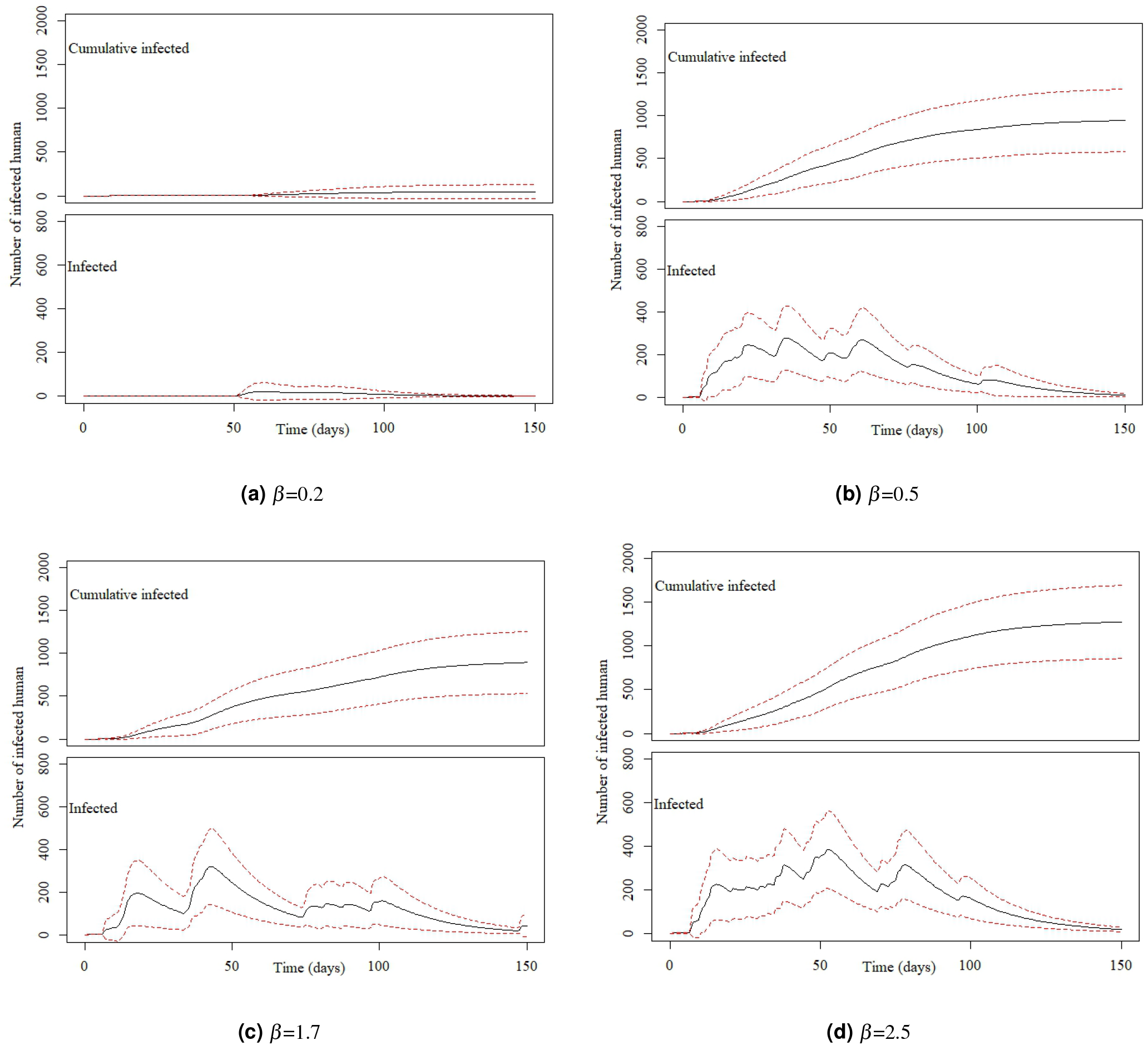
Number of infected individuals in the Uganda Ebola network with 95% confidence interval with *P*_0_=0.1, *γ*=0.5, and for (a) *β* =0.2, (b) *β* =0.5, (c) *β* =1.7, and (d) *β* = 2.5

Figure 8 shows risk maps for *P*_0_= 0.1 and *γ* = 0.5. It is evident from the map that, all districts in our network are at low risk for EVD spreading for selected values of *β*.

**Figure 8.**
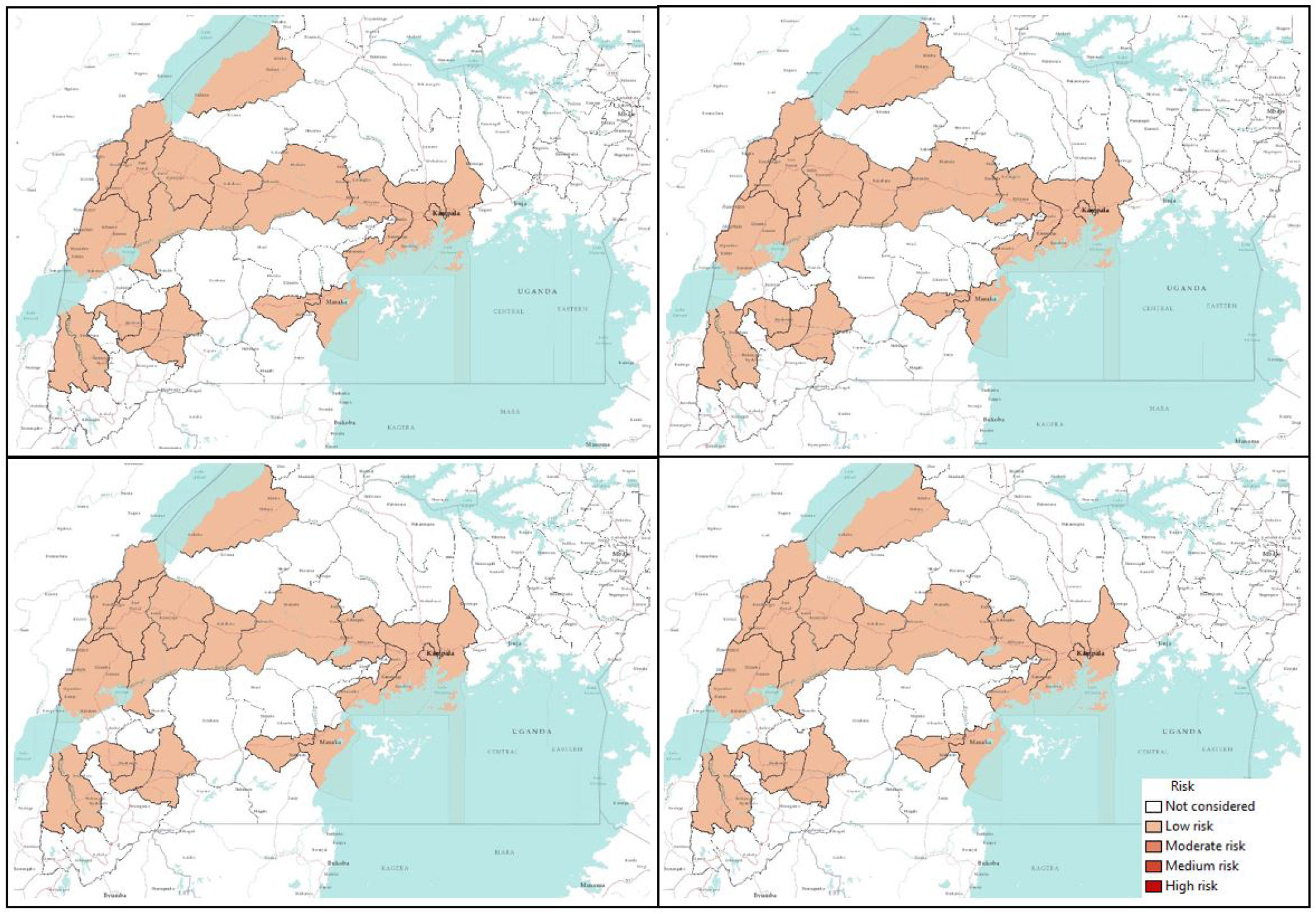
Risk map of Ebola spreading within selected 23 districts in Uganda for *β* =0.2 (top left), 0.5 (top right), 1.7 (bottom left), 2.5 (bottom right). The map is colour coded according to the risk of Ebola spreading.

Further decreasing the value of *γ* to 0.1 decreases cumulative number of infected humans as well as peak infection size. Figure 9 shows that, when *β* is 0.2 and 0.5, the EVD does not spread at all and stays within the outbreak location with only 2-3 infected persons. However, increasing the value of *β* increases number of infected humans but the cumulative number of infected remains less than 150 even for our highest value of transmission rate (*β* = 2.5).

**Figure 9.**
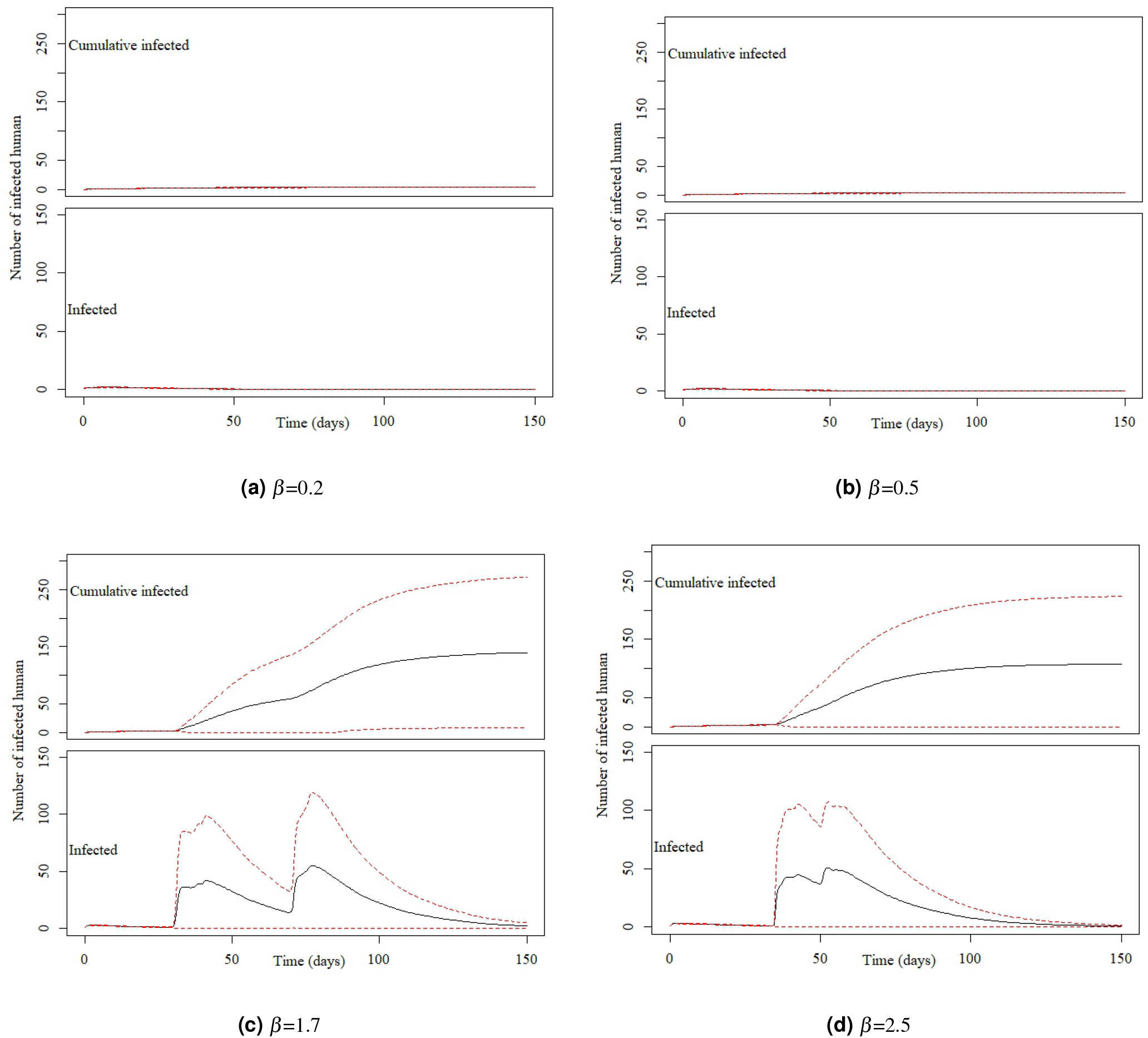
Number of infected individuals in the Uganda Ebola network with 95% confidence interval with *P*_0_=0.1, *γ*=0.1, and for (a) *β* =0.2, (b) *β* = 0.5, (c) *β* =1.7, and (d) *β* = 2.5

Figure 10 shows no risk of EVD spreading for *β* =0.2 and 0.5. However, with increasing *β*, all our districts are at low risk of EVD spreading.

**Figure 10.**
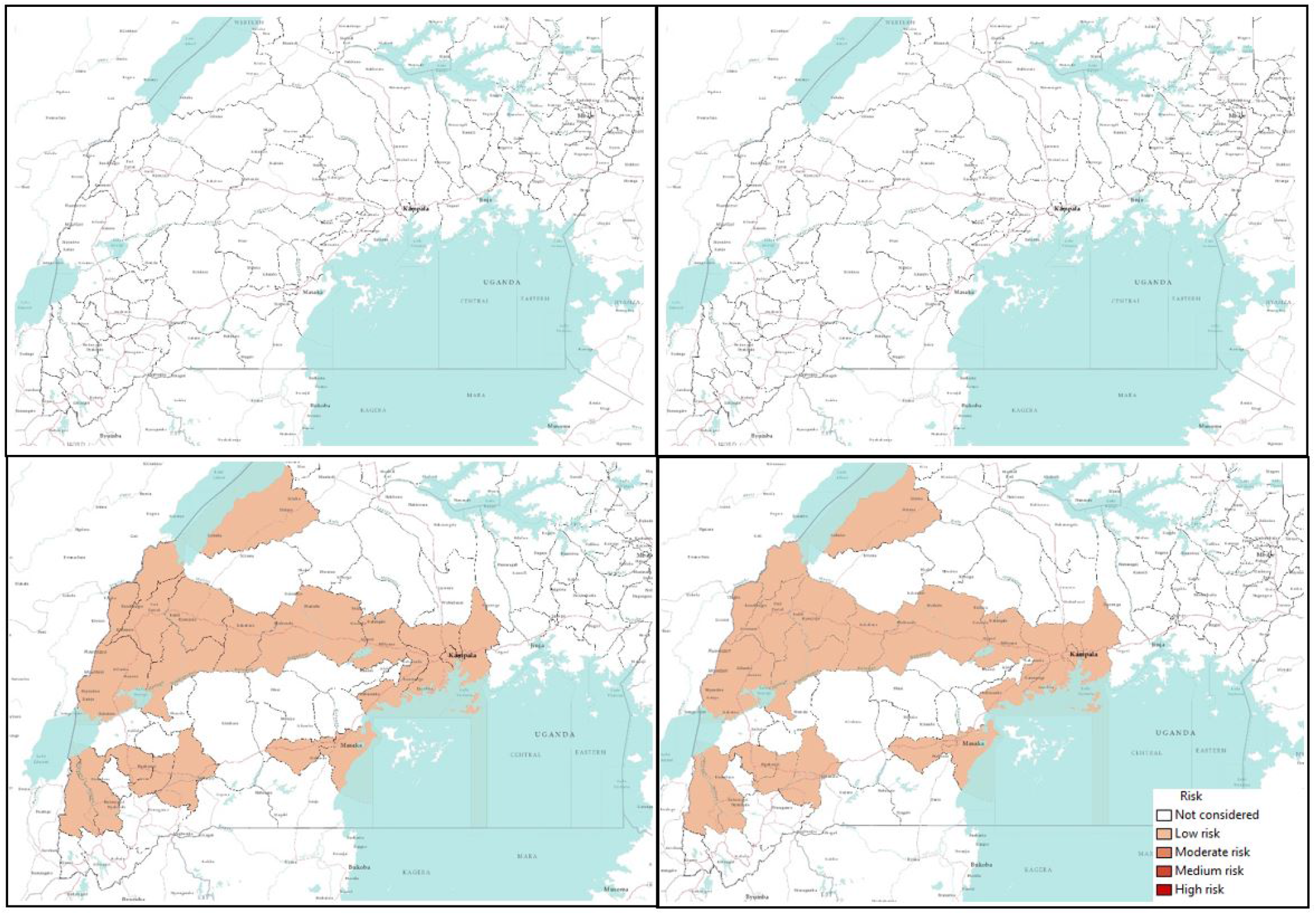
Risk map of Ebola spreading within selected 23 districts in Uganda for *β* =0.2 (top left), 0.5 (top right), 1.7 (bottom left), 2.5 (bottom right). The map is colour coded according to the risk of Ebola spreading.

Summarizing simulation results and risk maps presented in Figures 6-9, we can see a significant reduction in the epidemic size and simultaneously infected humans when *P*_0_=0.1. Therefore, a reduction in probability of pathogen transfer via a potential link i.e. reduced physical contact between humans while they are active/mobile greatly reduce the number of infected as well as severity of EVD. It also reduces the risk of EVD spreading.

EVD spread from physical contact with bodily fluid such as blood, feces, vomit, saliva, mucus, tears, breast milk, urine, semen, sweat etc from infected persons. Therefore, a susceptible person can only be infected with EVD if it comes in contact with these bodily fluids from an infected human. Therefore, if infected people are identified and moved to quarantine and restrict to travel, then the infection can be contained. However, non-hospitalized people during early symptomatic stage keep moving for their daily life and spread the infection with people they come in contact with. Early detection in countries where people are not very health conscious is very challenging. Therefore, when an infected case is found, if human movements are reduced significantly, the spread can be contained. Our simulation results conform with this movement reduction and reduced EVD infected cases. However, this not practical as human movements can not be controlled.

Activity rate (*γ*) can be translated to human movements within the network, therefore increasing activity rate means increased but shorter duration human movements. From all simulation results, it is evident that an increase in activity rate increases the epidemic size and as well as the speed of the infection to reach its peak. Therefore, increasing activity rate means a severe and fast EVD spreading. Simulation results suggest that, short duration but frequent human movements result in greater number of infected humans and high risk of EVD spreading than longer duration but less frequent movements.

The probability at which infection spreads to susceptible people via a potential contact (*P*_0_) has obvious impacts on the epidemic size. Increasing *P*_0_ increases the epidemic size irrespective of the activity rate *γ* which is evident from Figures 2,4,6, and 8. Comparing Figure 3 and Figure 7 as well as Figure 5 and Figure 9, for similar values of the activity rate, decreasing *P*_0_ decreases the number of infected humans significantly. *P*_0_ can be translated to the probability of physical contact of infected mobile humans with others. Therefore, our simulation results conforms with another mitigation strategy against EVD spreading which is to stay away from contact with people whose status with respect to EVD are not know. The lower value of *P*_0_ can be achieved by minimizing physical contact among people during movements/travels. If people are aware of the risk of EVD spreading in their area and keep themselves from physical contact with others, it will significantly reduce infected cases in case of an outbreak.

Risk maps shows that there some districts as shown in Table 2 and 3, who are at higher risk of Ebola spreading in our network for our specific network scenario. The risk assessment provides an important information for Ebola preparedness. Ebola preparedness includes setting up medical facilities as well as employing different preventive measures for Ebola spreading. However, resources are limited and it is always essential to find a way utilize the resources in a fruitful way. However, our risk only showed examples of of our developed risk assessment method where we have used very limited generalized movement data. Although proposed risk assessment method have the capability to provide guidelines for public health people, it requires more accurate movement data to be used practically in the field. Future work will include an evaluation of the impact of major EVD intervention pillars (i. e., coordination, vaccination, surveillance, risk communication, case management, and safe burials) on the risk assessment, to provide a realistic picture under multiple scenarios.

## Conclusions

We present a novel method for risk assessment based on a multilayer temporal network. This method has the ability to assess risk of EVD spreading when accurate network data is available and can be a important tool for public health people during an outbreak. We demonstrate an application of our developed method using a multilayer temporal network formulated with generic and incomplete movement data in Uganda. Simulations results from this multilayer temporal network confirm that reduced physical contacts with people while travelling, and taking other preventive measures decrease the risk of EVD spreading. Simulations also show that some districts are at higher risk than others in the considered scenario. This provides public health personnel directions about prioritizing their effort to limit EVD spreading during an outbreak. However, assessed risks are crucially dependent the network structure and can be fully trusted for resource allocation once accurate individual-level movement data in time and space are available.

## Acknowledgement

The authors gratefully acknowledge the financial support provided by United States Department of Agriculture under grant numbers 3020-32000-008-04-S and 2015-67013-23818.

## Acknowledgement

M.H.R and C.S formulated the model, M.H.R, C.S, and M.S analyzed the data, M.H.R and C.S developed the method, M.H.R. developed the simulation tool, M.H.R and C.S. analyzed the results and contributed to the discussion, M.H.R and C.S. prepared the first draft, M.S., F.O., and I. M. helped in revising and rewriting the draft, provided important feedback on the results. All authors reviewed the manuscript.

## Additional information

### Competing interests

The authors declare that they have no competing interests.

